# Ant impacts on global patterns of bird elevational diversity

**DOI:** 10.1101/2023.04.29.538805

**Authors:** Umesh Srinivasan, Kartik Shanker, Trevor D. Price

**Author notes:** Corresponding author: Umesh Srinivasan. **Author contributions:** US, KS and TDP co-conceived the study, US and TDP performed the analyses. US wrote the first draft and all authors contributed to revising the manuscript.

## Abstract

Across the world’s mountains, elevation-species richness relationships are highly variable. Here, using data on bird species elevational distributions from all 46 of the world’s major mountain ranges, bird species dietary traits, and the distribution of the low-elevation ant genus *Oecophylla*, we show that global patterns in bird elevational diversity are likely to be affected by competition with ants. *Oecophylla* is an exceptionally abundant and aggressive predator of invertebrates, which preys on the same species that sympatric invertivorous bird species feed on. In mountain ranges with *Oecophylla* present in the foothills, maximum species richness of invertivorous birds occurs, on average, at 960m, ∼450m higher than in mountain ranges without *Oecophylla*. Further, in mountain ranges with *Oecophylla*, species richness of invertivorous birds increases initially with with elevation to produce a mid-elevation peak in invertivore bird species richness. Where *Oecophylla* is absent, invertivore bird species richness generally shows monotonic declines with increasing elevation. We attribute the pattern to the following mechanism: first, *Oecophylla* reduces prey density for invertivorous birds; second, low invertebrate prey abundance reduces invertivorous bird density and third, lower bird density is correlated with lower bird species richness. Because invertivores dominate montane bird communities, global elevational bird diversity patterns are also driven by *Oecophylla*. The findings emphasize how competitive interactions between distantly related taxa set geographical range limits.

## Introduction

A large fraction of the world’s terrestrial species are found in mountainous regions, especially in the tropics (1) and the distribution of the elevational ranges of all species on a mountain generates an emergent relationship between elevation and species richness through range overlaps. Especially in the tropics, many montane species have small elevation ranges. Such species are likely to be thermal specialists (2, 3) and are consequently experiencing disproportionately high impacts of climate change (4), making it particularly important to understand why and how richness patterns vary with elevation. Mechanisms hypothesised to limit elevational ranges and therefore regulate species richness along elevational gradients include abiotic factors (e.g. temperature gradients and species’ thermal tolerances (2)), biotic interactions (e.g., interspecific competition, (5)) and geometrical factors (i.e., area available at different elevations, (6)).

Competition has been widely invoked as important in limiting elevational ranges, based on first, the distributions of closely related species, which are expected to be ecologically similar (5, 7) and second, negative relationships between regional species richness of a taxon and elevational range sizes (8). However, competition between distantly related taxa may affect population sizes and distributions even more dramatically than competition between related species if the larger functional differences between taxa reflect superior abilities in particular environments (9–12).

Species richness patterns vary greatly across elevational gradients; in some locations, more species are found at the mountain base rather than higher up, but in others the peak in species richness lies at intermediate elevations (6, 13). Here, using published datasets on the elevational distributions of bird species in the world’s mountains (8, 13; Fig. 1), we test the hypothesis that global patterns in the elevation-species richness relationship of montane birds are affected by competition across phyla, specifically between montane invertivorous birds and weaver ants (*Oecophylla*) – a low-elevation (up to ∼900m to 1000m ASL) genus of unusually aggressive, abundant and predatory invertivorous ants (14, 15). *Oecophylla* is represented by two tropical and sub-tropical allopatric species; *O. longinoda* occurs in sub-Saharan Africa (but not Madagascar), whereas *O. smaragdina* is found in south and southeast Asia, the Malay Archipelago and northern Australia. The genus is absent from the neotropics and temperate regions (14).

**Fig. 1.**
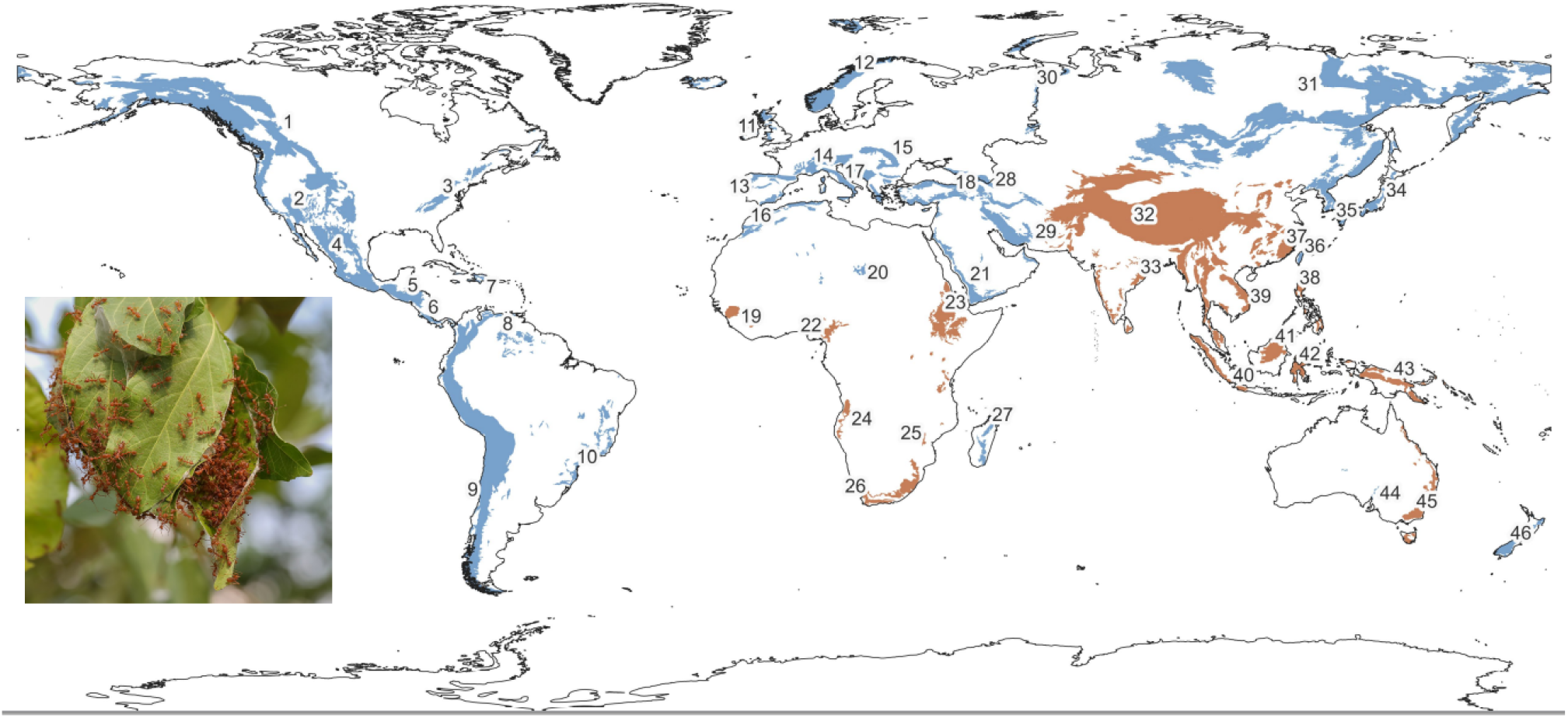
Map of the world’s 46 major mountain ranges (13), color coded based on whether species in the ant genus *Oecophylla* occurs in the foothills (orange) or not (blue). The inset shows an arboreal nest of *Oecophylla smaragdina*, the Asia-Pacific representative of the genus. (photograph: Basile Morin/Wikimedia Commons). For a list of the mountain ranges corresponding to the numbers on the map, see Table S2.

*Oecophylla* species are highly effective arthropod predators and are consequently widely used for biological control (16, 17). Indeed, *O. smaragdina* was the first known biological control agent, dating to 304 CE in China (17, 18). Experimental manipulations have found that the presence of *Oecophylla* generates substantial (∼15% on average) reductions in both pests and pest damage (*N* = 25 studies, 16). In a mango orchard in Australia, Pinkalski et al. (19) found that *Oecophylla* occupied about half the trees at any one time, with an average of at least 30,000 workers on each tree, and surveys suggested that these abundances are very likely similar to those found in adjacent natural forest habitats. In mangroves, leaf damage by arthropods was found to be four times higher on trees without *Oecophylla* than with *Oecophylla* (20). In a subtropical lowland forest in the eastern Himalaya, the removal of *O. smaragdina* generated a threefold increase in the abundance of non-ant arthropods over one month, especially in the orders Lepidoptera and Coleoptera, the two orders that form primary components of the diets of sympatric insectivorous bird species (21).

If *Oecophylla* reduces prey density for invertivorous birds via exploitation competition, the presence of this ant should also reduce the number of invertivorous birds at low elevations. A reduction in the abundance of invertivorous birds should then lead to a reduction in species richness, because lower abundances of individual species increases their risk of extirpation, thereby reducing long-term persistence (22–24). Empirically, both the positive correlation between resource density and consumer density, and between consumer density and species richness has been demonstrated for invertivorous birds across elevational gradients (25–27). More direct associations between arthropod abundance and species richness have also been demonstrated (26–29).

Based on this reasoning and the global distribution of *Oecophylla*, we predicted that (1) in mountain ranges where an *Oecophylla* species is present in the foothills, the peak in species richness of invertivorous birds would be at higher elevations than it would be in mountains where *Oecophylla* is absent, (2) the higher elevation richness peak in *Oecophllya* locations would hold true for omnivorous bird species, although the strength of these patterns would be weaker than for invertivores because omnivores should compete less with *Oecophylla*, (3) other avian dietary guilds (such as nectarivores, scavengers), which are not expected to compete with *Oecophylla*, should show no consistent association with *Oecophylla* presence/absence. Following from the above predictions, we expected that (4) for invertivores—and to a lesser extent for omnivores—the presence of *Oecophylla* in mountain foothills would result initially in a positive relationship between elevation and species richness until a mid-elevation peak, after which species richness would decline; for non-invertivores, this initial relationship would not be impacted by the presence/absence of *Oecophylla*, and would be negative or “flat” across of the world’s mountains.

Using published elevational ranges from 46 of the world’s major mountain ranges (Fig. 1, (13)) we compared the average elevation at which species richness peaks for the three avian dietary guilds (invertivores, omnivores and all others combined) in the presence and absence of *Oecophylla*. In addition to *Oecophylla* as a predictor variable in a linear model, we also tested whether net primary productivity and precipitation peaked at the same elevations as species richness, because both have been postulated to be important determinants of bird species elevational patterns (6, 30). Because there might not be clear peaks in species richness for particular diet guilds in particular mountain ranges, we used a second method to test our predictions. We used automated piecewise regression to test whether, at low elevations, the relationship between elevation and species richness for these dietary guilds was positive, negative, or “flat” (see Materials and Methods; Fig. 2).

**Fig. 2.**
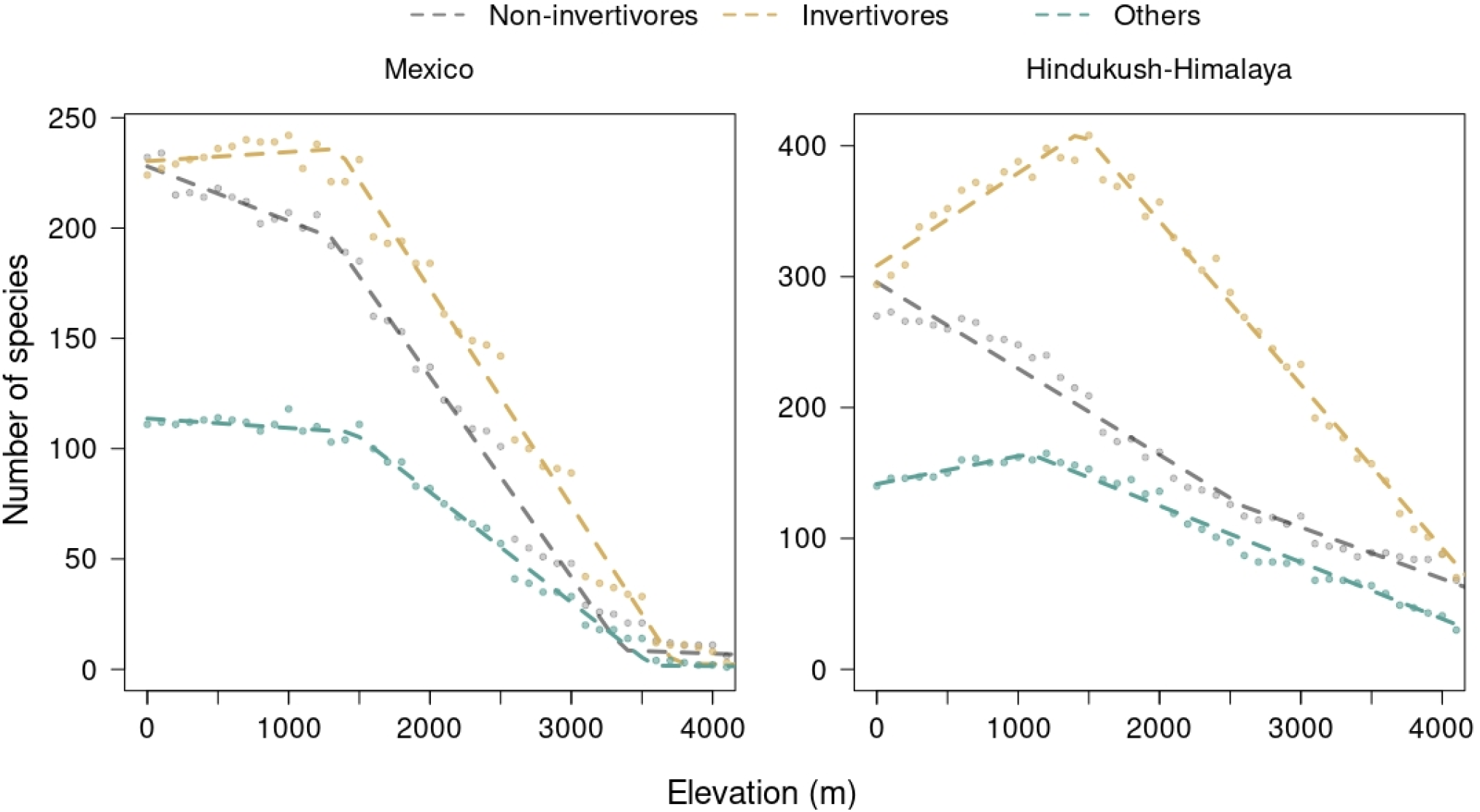
The relationship between species richness and elevation in two example mountain ranges, one lacking *Oecophylla* in the foothills (Mexico) and one with *Oecophylla* at low elevations (Himalaya). Dashed lines show fits from an automated piecewise (or segmented) regression performed using the *segmented* package (55) in R (56). At low elevations, the relationship between invertivore and omnivore species richness can be either positive (Himalaya), flat (Mexico) or even negative (e.g., see UK and Ireland, Northern South America in Fig. S4). In general, others show monotonic declines with increasing elevation (Fig. S4). See Table S2 for estimated slopes and 95% confidence intervals of the first segment of the piecewise regression for different diet guilds in different mountain ranges.

## Results

Overall, a linear model with *Oecophylla* presence/absence, bird dietary guild, productivity and precipitation explained ∼47% of the variability in the elevation at which bird species richness reaches its maximum across the world’s mountain ranges (*R*^*2*^ = 0.47). Only diet and the presence or absence of *Oecophylla* contributed significantly to the model (diet: *F*_2,126_= 33.07, *p* < 0.01; *Oecophylla*: *F*_1,126_ = 40.85, *p* < 0.01). Across the world’s mountain ranges, the mean elevation at which invertivore richness peaks in the presence of *Oecophylla* is 960m ASL ± 62m SE, over 400m higher than in mountains without *Oecophylla* (535m ± 50m; Fig. 3). This pattern was similar but weaker for omnivores (*Oecophylla* present: 710m ± 91m; *Oecophylla* absent: 382m ± 62m; Fig. 3). The species richness of other guilds combined generally reaches a maximum at low elevations (well within the elevational range of *Oecophylla*: 77m ASL ± 30m SE in the presence of the genus and 367 ± 102m SE in its absence; Fig. 3). Within any dietary guild, elevations at which productivity and rainfall reach their maximum were unrelated to the elevations at which bird species richness peaks (productivity: *F*_1,126_ = 0.59, *p* = 0.44; precipitation: *F*_1,126_ = 0.64, *p* = 0.43; productivity-diet interaction: *F*_1,126_ = 0.02, *p* = 0.99; precipitation-diet interaction: *F*_1,126_ = 0.22, *p* = 0.80; Fig. S1).

**Fig. 3.**
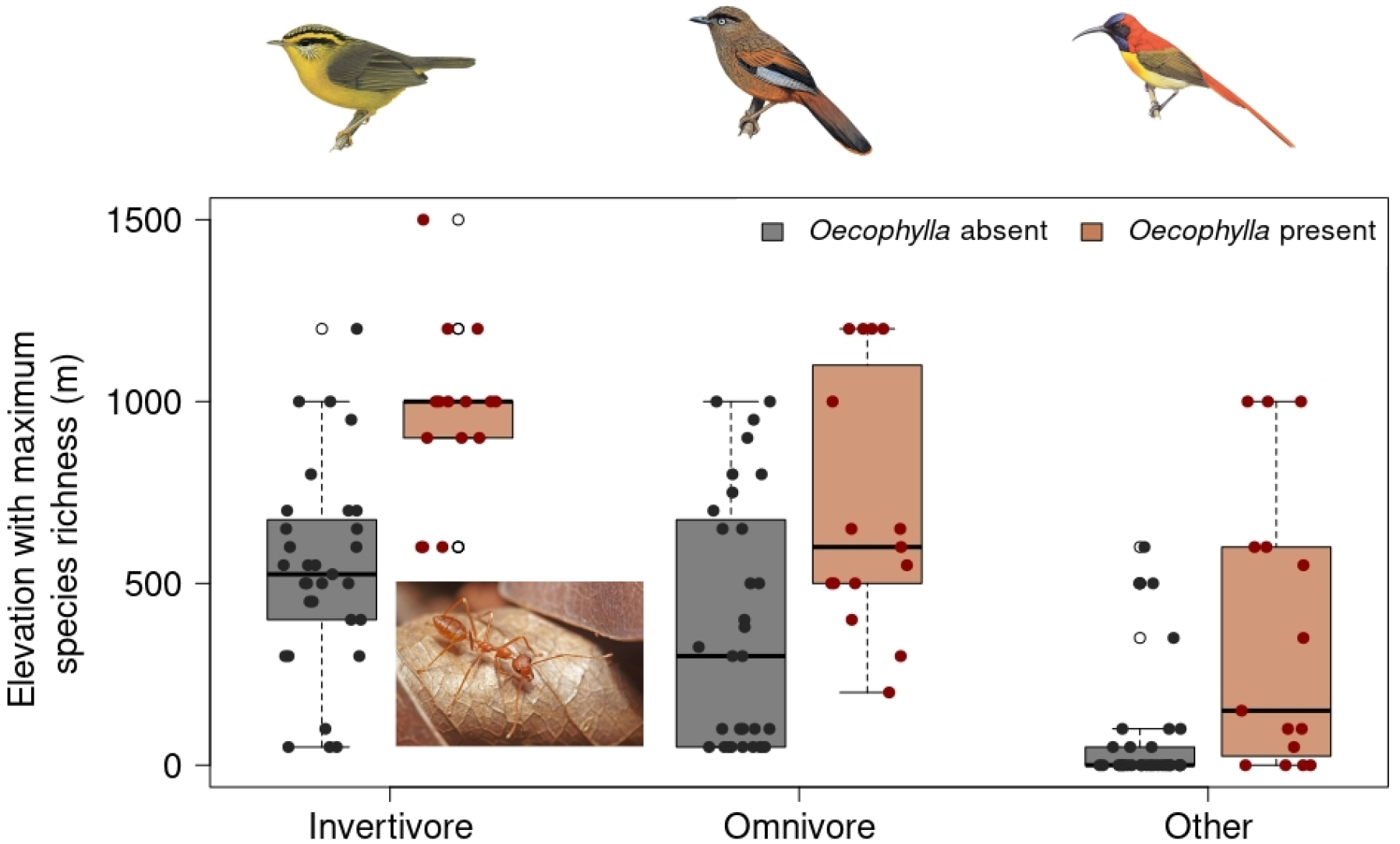
Boxplot of the relationship between the presence/absence of the invertivorous ant genus *Oecophylla* in the foothills of the world’s mountain ranges and elevation with the maximum species richness of invertivorous, omnivorous and other guilds combined (mean peak richness elevation across three dietary guilds. In the boxplots, solid horizontal lines represent the medians. Each point represents one of the world’s 46 major mountain ranges. Overlaid dots represent individual mountain ranges, and are jittered for clarity. The photograph is of *O. smaragdina*, the Asia-Pacific representative of the genus (photograph: Muhammad Mahdi Karim/Wikimedia Commons). Bird images (from left to right, Yellow-throated Fulvetta (*Schoeniparus cinereus*), Blue-winged Laughingthrush (*Trochalopteron squamatum*) and Fire-tailed Sunbird (*Aethopyga ignicauda*) are courtesy Birds of the World, and represent each of the diet guilds.

Restricting our analysis to tropical mountains, we found similarly strong patterns for invertivores, slightly lower associations for omnivores and no association for other guilds combined (Fig. S2). (See also Table S1 for an ANOVA table from a multiple regression where 10 diet guilds were considered rather than three.) Because invertivores and omnivores tend to dominate montane bird communities (60% ± 7% SD of the species of montane bird communities are invertivores/omnivores; (31)), total species richness follows the invertivore pattern (31).

We conducted two additional analyses to test these findings. First, we obtained similar results when we analysed a different dataset on elevational ranges, collected from 31 mountain ranges (eight with *Oecophylla* at their base, 23 without *Oecophylla*) using citizen science (8, Fig. S3). From this same dataset we also found that elevations of mountain bases did not differ significantly between mountain ranges with and without *Oecophylla* (b_with *Oecophylla*_ = 20.11 ± 27.67m higher; *R*^2^ = 0.02; one-way ANOVA). Second, to the level of resolution attainable, species richness often plateaus across elevations, especially at mid to low elevations (6), with the result that peak positions can be uncertain (examples in Fig. S4 include all guilds in the Sahara, invertivores and omnivores in Madagascar, Mexico, Zagros and other ranges). We used automated piecewise regression to test whether, at low elevations, the relationship between elevation and species richness for these dietary guilds was positive, negative, or “flat” (i.e., statistically not distinguishable from zero; Fig. 2). In mountain ranges with *Oecophylla* at low elevations, species richness increases with increasing elevation for invertivores and omnivores (i.e., declines towards the mountain base from a mid-elevation hump), whereas this relationship is either “flat” (i.e., statistically not distinguishable from zero) or negative in mountains without *Oecophylla* (Fig. 4, Table S2). The presence or absence of *Oecophylla* is unrelated to low-elevation species richness patterns for other dietary guilds combined (Fig. 4, Table S2).

**Fig. 4.**
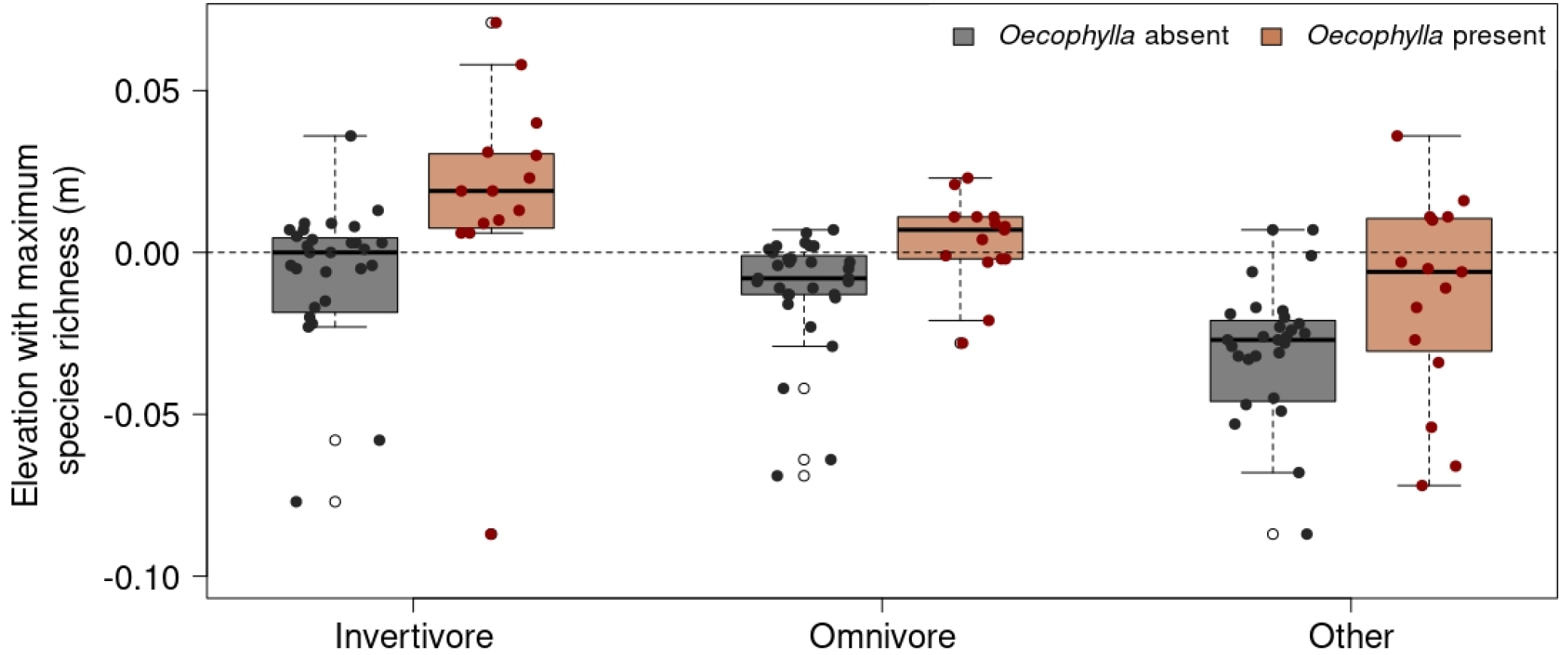
Boxplots of the slope of the first segment (at low elevations) of piecewise regressions relating species richness to elevation (see Fig. 2) for different bird diet guilds in relation to the presence/absence of *Oecophylla* ants at mountain foothills. For invertivores in mountains where *Oecophylla* is present, the slope of the first segment is always positive, consistent with lower species richness at mountain bases and a mid-elevation peak in species richness. Where *Oecophylla* is absent, this slope is statistically indistinguishable from zero (i.e., constant species richness with increase in elevation followed by a decline; see Mexico in Fig. 2) or negative (i.e., species richness declines with increasing elevation from the mountain base). This pattern is present also for omnivores but weaker. For non-invertivores, the slope is not different between mountains where *Oecophylla* is present or absent and is typically negative.

## Discussion

The presence or absence of *Oecophylla* ants in tropical and sub-tropical lowlands is an important predictor of the elevation at which avian invertivores are most diverse. Previously, a number of factors have been hypothesized to drive species richness patterns across elevations, including the distribution of net primary productivity (6, 30), precipitation (32, 33) and mountain shape (i.e., the extent to which different elevational belts differ in area (6, 34). When all regions are combined, we found little evidence that either productivity or precipitation are important determinants of elevational species richness patterns (Fig. S1). However, some mountain ranges with arid bases and greater precipitation at higher elevations have bird species peaks at mid-elevations, likely attributable to higher productivity and structural diversity (6). Relationships between species richness and productivity/precipitation might hold for certain taxa and mountain ranges; however, our results (Fig. S1) indicate that at the global scale, and for birds, these patterns are not general. Instead, we find the presence of *Oecophylla* provides a stronger signal in the data.

It is conceivable that large areas in the lowland tropics have led to the production and maintenance of many species, an “age and area” hypothesis (35). This may then lead to a linear decline in species richness with elevation, for instance on the east flank of the Andes (36). Nevertheless, neotropical mountains adjacent to smaller tropical lowlands and temperate areas also show linear declines. Indeed, previous tests of bird elevational diversity within mountain ranges have rejected the importance of area (6). More recently, Elsen and Tingley (34) classified mountains into four shapes (a) diamond, with maximum area at mid-elevations, (b) pyramid, with maximum area at low elevations (c) inverse pyramid, with maximum area at high elevations, and (d) hourglass, where mountains have a mid-elevation increase in area but maximum area occurs at high elevations. We found no systematic differences in the frequency of these mountain types between regions with and without *Oecophylla* (see Supplementary Table 3 in (34)).

Large impacts of ants on tropical vertebrates at low elevations are highly likely given that the global biomass of ants is higher than that of all wild birds and mammals combined (37). Approximately one-third of this biomass is found in low-elevation tropical forests (37). *Oecophylla* densities alone have been estimated at 250 ants m^−2^ (19), suggesting a likely large impact of these predatory ants on arthropods with ramifying impacts on other arthropod-consuming species. Indeed in *Oecophylla* regions, ant predation has been experimentally demonstrated to be the major cause of insect mortality in the lowlands (38, 39). In these regions, ant diversity steeply declines with elevation, and few ants are found in mid-elevations (1500m-2500m, 40, 41), freeing up resources for birds (21). In New Guinea, invertivorous birds have been shown to be the main predator on arthropods at mid-elevations (39)

The decline in ant abundance with elevation applies also in tropical locations without *Oecophylla* (42, 43). However, in these regions, bird species richness patterns often show monotonic declines with elevation (Fig. S4). Although monotonic declines may reflect climate (e.g., linear declines in productivity), predatory ants are present in the neotropics (44), and if they were to reduce arthropod populations to a similar extent as *Oecophylla*, might be expected to lead to mid-elevation peaks in insectivorous birds. However, ants can be both competitors and facilitators. Unlike in the palaeotropics, in the low-elevation neotropics, army ants (*Eciton, Labidus* sp.) are common and consume terrestrial invertebrates and small vertebrates, but unlike *Oecophylla*, they facilitate the foraging of hundreds of invertivorous bird species, rather than competing with them. At least 465 species of lowland neotropical birds have been recorded following army ants; 50 of these species are “professional ant followers”, relying entirely or mainly on ants to find food (45). Indeed, obligate ant followers are parasites on army ant swarms, “stealing” about 30% of the daily leaf litter food requirement from migrating ant colonies (46). Thus, while army ants and low-elevation invertivorous neotropical birds might be engaged in exploitation competition (because of yet-to-be investigated dietary overlaps), army ants are still indispensable facilitators for dozens of bird species in obtaining food on a daily basis (47).

The spatial pattern we have identified is similar to that associated with temporal turnover in Europe roughly 34 million years ago. Fossil evidence from Europe shows that there were many more ant species belonging to tropical genera, including several species of *Oecophylla*, during the relatively warm Eocene (48). In the cooler Oligocene, these ants disappeared and were replaced by songbirds (Passeriformes: oscines) (49, 50). Especially in the current climate crisis, such historical patterns coupled with our findings emphasize the need to consider how species interactions across phyla might set geographical range limits. Finally, our results demonstrate the great dangers in introducing *Oecophylla* for biocontrol to locations where the genus is presently absent, such as the neotropics.

## Materials and Methods

Data on the elevational ranges of the world’s montane birds were taken from supplementary materials to two publications (8, 13). The paper by (13) was based on published elevational ranges, the majority of which come from (51). Ranges were assigned to all 46 of the world’s major mountain ranges (as defined by the Global Mountain Biodiversity Assessment; https://www.gmba.unibe.ch/research/). On the other hand Freeman et al. (8) used different criteria to delineate 37 mountain ranges, from which they analysed citizen science data from 31 (African mountain ranges are entirely absent from the data they analyse). Because the two sources (a) use different criteria to demarcate different mountain ranges, and (b) different sources of elevational range data for birds, we tested whether our results were concordant across the two datasets. We present the data from ref (13) as our primary dataset for analysis, because of its higher geographical coverage of the world’s mountains, and its emphasis on the distribution of richness of all species along elevation gradients.

We first used a linear model to ask whether the elevations at which invertivorous bird species richness reached their maximum was related to the presence of *Oecophylla* in the foothills. We also tested whether estimates of precipitation (52), and of net primary productivity as it would be in the absence of humans (53), peaked at the same elevations as species richness, because both precipitation and productivity have been postulated to be important determinants of bird species elevational patterns (6). Our model was formulated as follows:

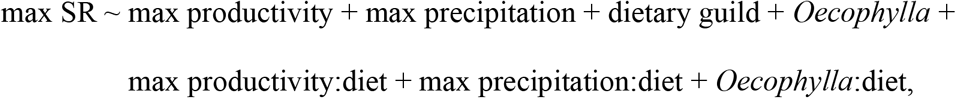

where:

a. max SR is the elevation at which the species richness of avian dietary guilds peaks across 46 mountain ranges (continuous variable),
b. max productivity is the elevation at which net primary productivity peaks (continuous variable),
c. max precipitation is the elevation at which precipitation peaks (continuous variable),
d. dietary guild is a factor variable with three levels (invertivore, omnivore and other; although a supplementary analysis included all ten dietary guilds instead of collapsing non-inverivore and non-omnivore diets into a single “other” category; see Table S1) (54) and
e. *Oecophylla* is a factor variable with two levels (*Oecophylla* present or absent in mountain foothills) (14). Interaction terms are indicated by the colon (:).

Using the dataset of (13), we conducted piecewise or segmented linear regression in the *segmented* package (55) in R (56). The *segmented* package allows for the detection of breakpoints where the slope in the relationship between two variables changes. In our case, for each mountain range in the dataset, the dependent variable was species richness (of invertivores, omnivores and other guilds combined, separately), and the predictor variable was elevation (Fig. 2). We constrained the regression to identify two breakpoints: (a) that differentiating elevation-species richness relationships between low and mid elevations, and (b) that separating elevation-species richness relationships between mid and high elevations. We did not specify any *a priori* expectations of where mid or high elevations started, allowing the model to select these automatically (Fig. 2). We selected these two breakpoints because (a) at low elevations, elevation-species richness relationships could either decline, remain unchanged or increase (in the case of a mid-elevation peak), (b) from mid to high elevations, the elevation-species richness relationships always declines (Fig. S4), and (c) at higher elevations, beyond which bird life is absent (especially in the highest elevation mountains such as the Hindukush-Himalaya and the Andes) or at mountain summits, the elevation-species richness relationship would switch to “flat” (i.e., elevations beyond which species richness is zero; see Fig. S4).

## Acknowledgments

We thank Ignacio Quintero for providing us with the shapefile of the world’s 46 major mountain ranges, Kavita Isvaran for inputs on the analyses, and Daniel Hooper and K. Supriya for inspiration.

