## Supplementary Information for "Ant impacts on global patterns of bird elevational diversity"

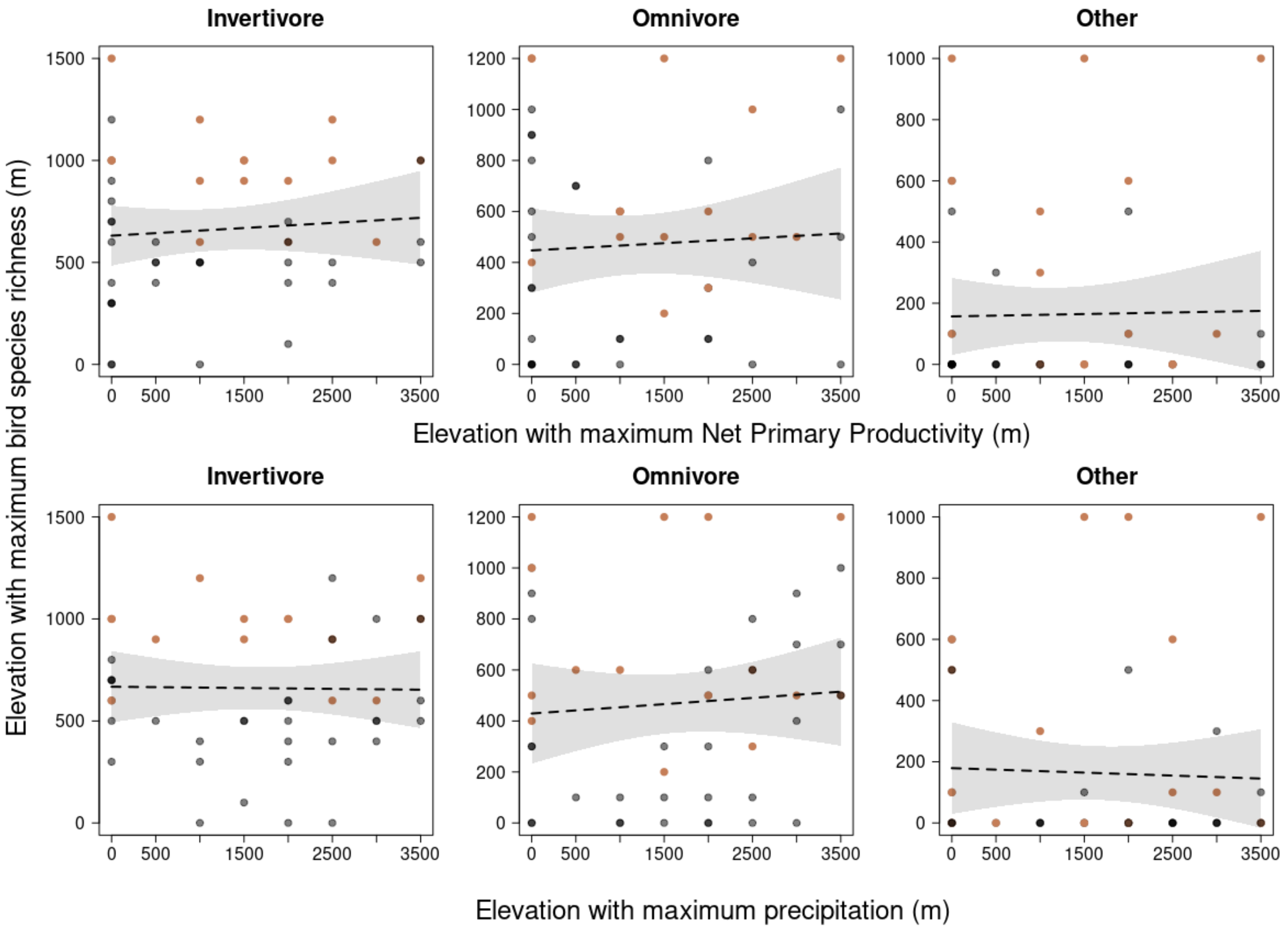


**Fig. S1. Across the world's 46 major mountain ranges, there is no evidence of a positive relationship between the elevations at which productivity or rainfall reaches a maximum, and the maximum point of species richness. Gray circles represent mountain ranges where the ant genus *Oecophylla* is absent in the foothills, and orange circles where the genus is present. Dashed black lines represent best fit linear regression estimates and gray polygons the standard error of the fit.**

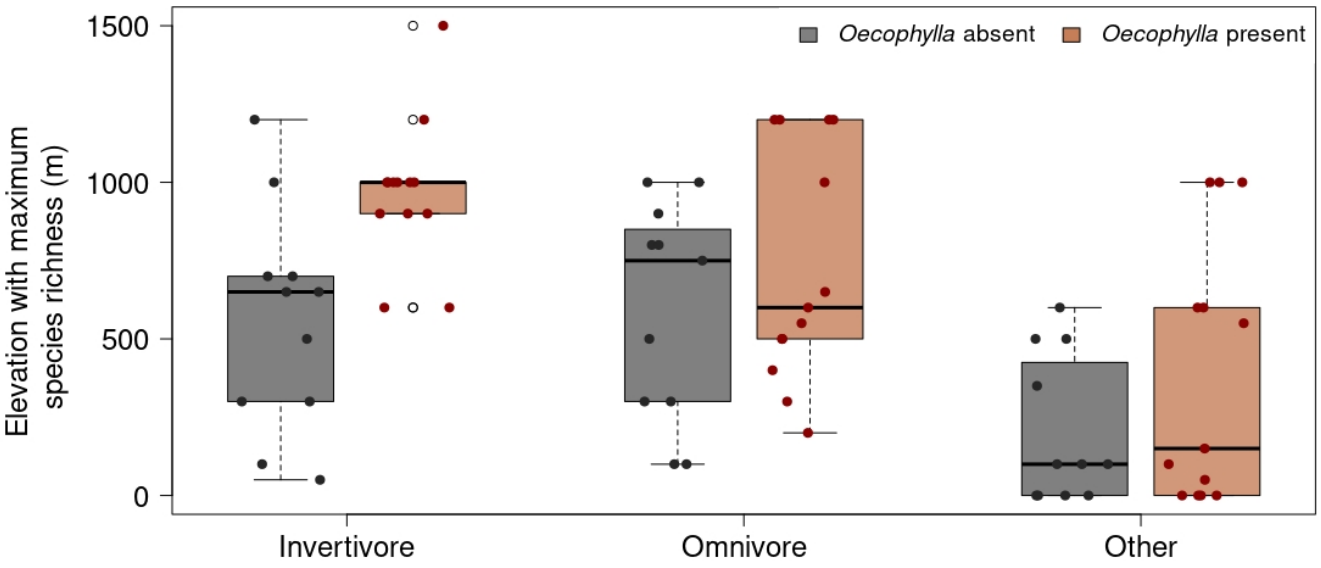


**Fig. S2. Boxplot of the relationship between the presence/absence of the invertivorous ant genus *Oecophylla* in the foothills of 24 of the world's tropical mountain ranges (present in 13 and absent in 11; marked with * in Table S2) and elevation with the maximum species richness of invertivorous, omnivorous and all non-invertivorous birds. On average, species richness of invertivores peaks 969 ± 63m SE in the presence of *Oecophylla* versus 559 ± 107m SE in its absence; the same comparison for omnivores is 730 ± 104m SE versus 595 ± 105m SE, and of non-invertivores 388 ± 116m SE versus 204 ± 71m SE.**

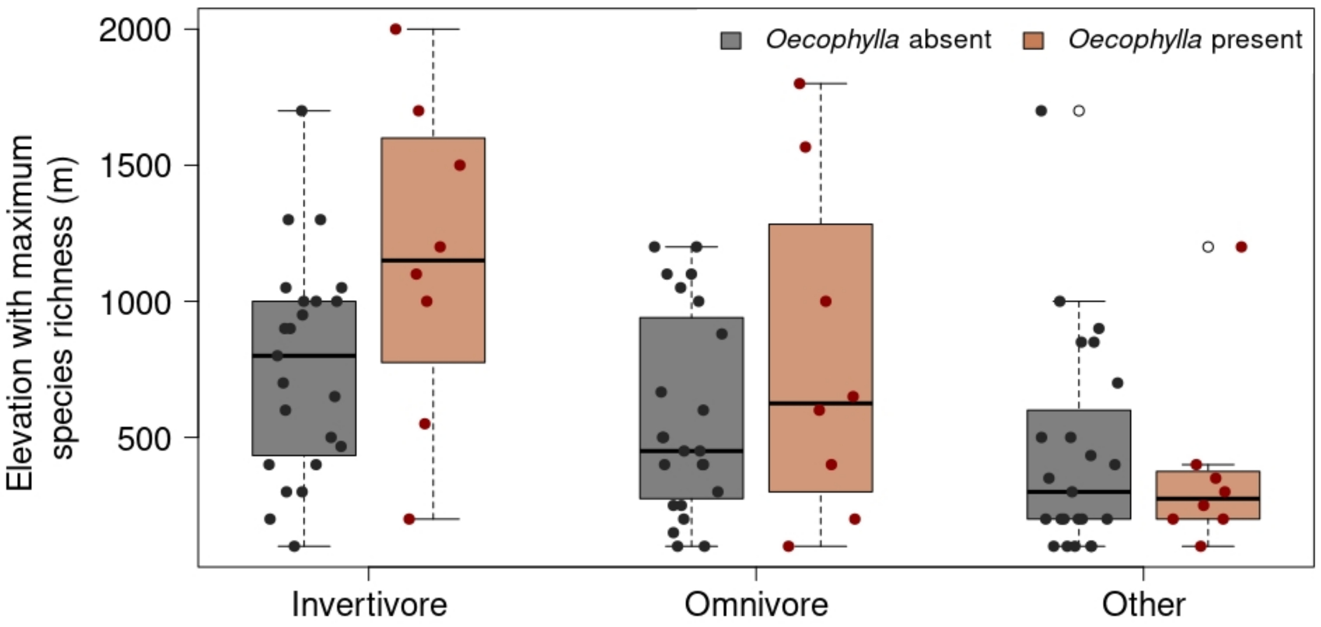


**Fig. S3. Boxplot of the relationship between the presence/absence of the invertivorous ant genus *Oecophylla* in the foothills of 31 of the world's mountain ranges (1) and elevation with the maximum species richness of invertivorous, omnivorous and others. On average, species richness of invertivores peaks 392 ± 94m SE higher, omnivores 213 ± 182m higher, and that of non-invertivores 68 ± 158m *lower* in mountain ranges with *Oecophylla* at low elevations.**

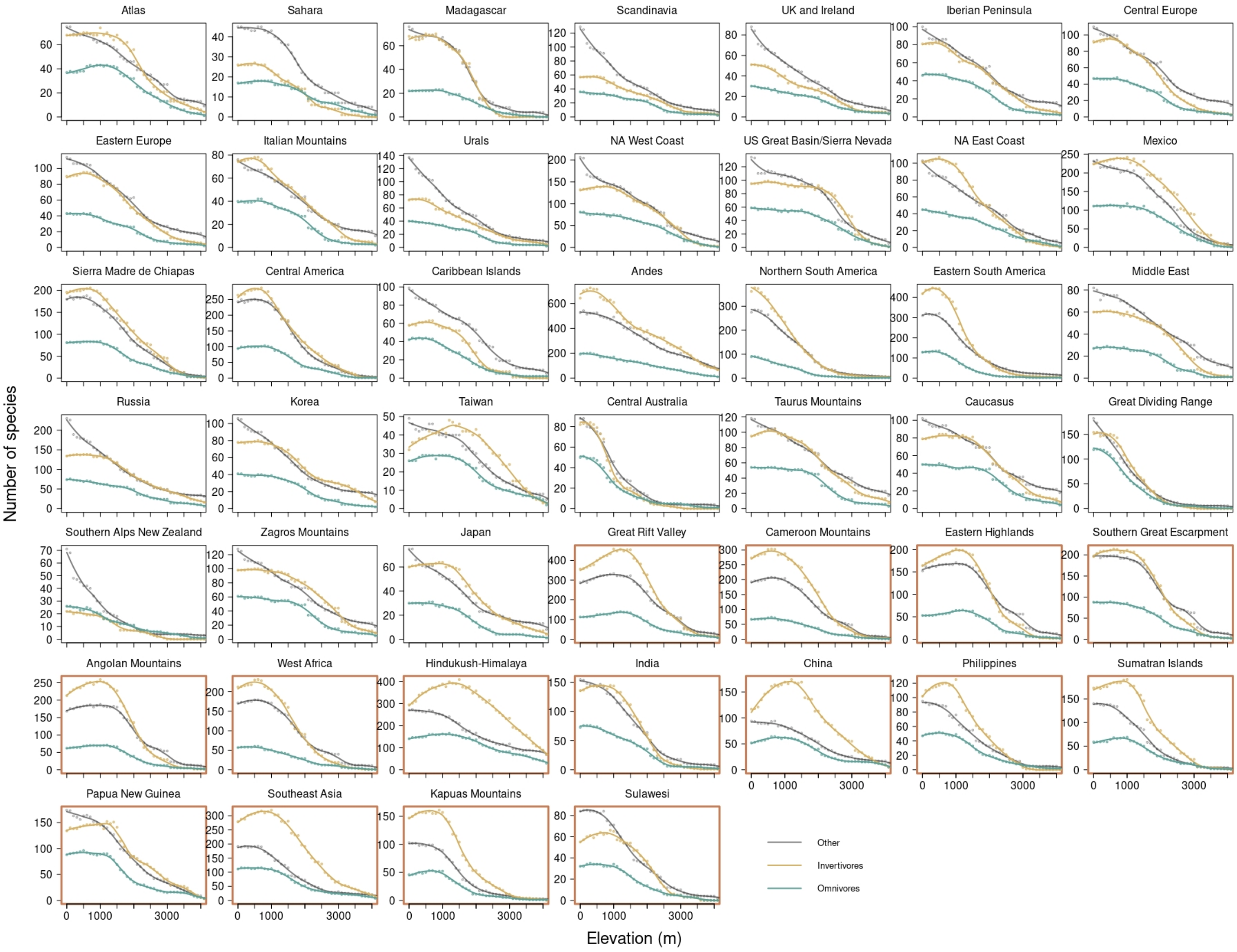


**Fig. S4. The relationship between elevation and species richness for invertivores (brown), omnivores (green) and non-invertivores (black) across the 46 major mountain ranges of the world (2). Curves are relationships estimated by locally estimated scatterplot smoothing (LOESS) in R (3). In general, the species richness of non-invertivores declines monotonically across all mountains, while that of invertivores and omnivores either declines monotonically or shows a mid-elevation peak. Species richness panels for mountain ranges where *Oecophylla* is present in the foothills are outlined in orange. Note that elevation is truncated to a maximum of 4,000m in the plots.**

**Table S1. Summary of Type III sum of squares from a multiple regression examining the role of net primary productivity, precipitation, avian dietary guild and the presence/absence at low elevations of species of the ant genus *Oecophylla* on the elevation at which bird species reaches reaches its maximum across the world's major mountain ranges, where 10 different dietary guilds are considered (4).**

| **Predictor variable** | **Sum of squares** | ***F*** | **df** | ***P*** |
|---|---|---|---|---|
| Intercept | 332 | 0.00 | 1 | 0.98 |
| Productivity | 1427 | 0.00 | 1 | 0.96 |
| Precipitation | 3 | 0.00 | 1 | 0.99 |
| Diet guild | 192501593 | 30.67 | 9 | <0.01 |
| *Oecophylla* presence | 4926 | 0.01 | 1 | 0.93 |
| Productivity : diet | 5697905 | 0.90 | 9 | 0.53 |
| Precipitation : diet | 5699618 | 0.90 | 9 | 0.53 |
| *Oecophylla*: diet | 32590426 | 5.12 | 9 | <0.01 |
| Residuals | 296991416 | - | 420 | - |
| **Total** | 533487646 | - | 460 | - |

**Table S2 | Slopes of the first segment of piecewise regressions (see Materials and Methods and Fig. S4) for different bird diet guilds in the 46 major mountain ranges of the world (2).**

| | | | Invertivores | | | Omnivores | | | Others | | |
|---|---|---|---|---|---|---|---|---|---|---|---|
| ID | Mountain Range | *Oecophylla* | Slope | Lower CI | Upper CI | Slope | Lower CI | Upper CI | Slope | Lower CI | Upper CI |
| 1 | NA West Coast | Absent | 0 | 0 | 0.01 | -0.01 | -0.01 | -0.01 | -0.05 | -0.05 | -0.05 |
| 2 | US Great Basin/Sierra Nevada | Absent | -0.01 | -0.01 | 0 | 0 | -0.01 | 0 | -0.02 | -0.02 | -0.01 |
| 3 | NA East Coast | Absent | 0.01 | 0 | 0.02 | -0.01 | -0.01 | -0.01 | -0.03 | -0.03 | -0.03 |
| 4 | Mexico* | Absent | 0 | 0 | 0.01 | 0 | -0.01 | 0 | -0.03 | -0.03 | -0.02 |
| 5 | Sierra Madre de Chiapas* | Absent | 0.01 | 0 | 0.03 | 0 | 0 | 0.01 | 0.01 | -0.01 | 0.03 |
| 6 | Central America* | Absent | 0.04 | 0 | 0.07 | 0.01 | -0.01 | 0.02 | 0.01 | -0.02 | 0.03 |
| 7 | Caribbean Islands* | Absent | -0.01 | -0.01 | 0 | 0 | 0 | 0.01 | -0.03 | -0.03 | -0.03 |
| 8 | Northern South America* | Absent | -0.18 | -0.18 | -0.17 | -0.04 | -0.04 | -0.04 | -0.12 | -0.12 | -0.12 |
| 9 | Andes* | Absent | -0.17 | -0.18 | -0.17 | -0.02 | -0.05 | 0 | -0.14 | -0.14 | -0.13 |
| 10 | Eastern South America* | Absent | -0.22 | -0.23 | -0.2 | -0.07 | -0.07 | -0.07 | -0.14 | -0.14 | -0.13 |
| 11 | UK and Ireland | Absent | -0.02 | -0.02 | -0.01 | -0.01 | -0.01 | -0.01 | -0.02 | -0.02 | -0.02 |
| 12 | Scandinavia | Absent | -0.02 | -0.02 | -0.02 | -0.01 | -0.01 | -0.01 | -0.05 | -0.06 | -0.05 |
| 13 | Iberian Peninsula | Absent | 0.01 | -0.01 | 0.01 | -0.02 | -0.02 | -0.02 | -0.03 | -0.03 | -0.03 |
| 14 | Central Europe | Absent | 0.01 | 0 | 0.01 | 0 | -0.01 | 0 | -0.03 | -0.03 | -0.03 |
| 15 | Eastern Europe | Absent | 0.01 | 0 | 0.02 | -0.01 | -0.01 | -0.01 | -0.03 | -0.03 | -0.03 |
| 16 | Atlas | Absent | -0.02 | -0.02 | -0.02 | 0.01 | 0.01 | 0.01 | -0.02 | -0.02 | -0.02 |
| 17 | Italian Mountains | Absent | -0.02 | -0.02 | -0.02 | -0.01 | -0.01 | -0.01 | -0.02 | -0.02 | -0.02 |
| 18 | Taurus Mountains | Absent | 0.01 | 0 | 0.02 | 0 | -0.01 | 0 | -0.03 | -0.03 | -0.03 |
| 19 | West Africa* | Present | 0.03 | 0.01 | 0.05 | 0 | -0.01 | 0.01 | 0.01 | -0.01 | 0.03 |
| 20 | Sahara* | Absent | 0 | 0 | 0 | 0 | 0 | 0 | 0 | -0.01 | 0 |
| 21 | Middle East | Absent | -0.01 | -0.01 | -0.01 | 0 | 0 | 0 | -0.02 | -0.02 | -0.02 |
| 22 | Cameroon Mountains* | Present | 0.02 | 0 | 0.04 | 0.01 | 0 | 0.02 | 0.02 | 0 | 0.03 |
| 23 | Great Rift Valley* | Present | -0.13 | -0.14 | -0.11 | 0.02 | 0.02 | 0.03 | 0.04 | 0.02 | 0.05 |
| 24 | Angolan Mountains* | Present | -0.09 | -0.1 | -0.08 | 0.01 | 0 | 0.01 | 0.01 | 0 | 0.02 |
| 25 | Eastern Highlands* | Present | 0.03 | 0.02 | 0.04 | 0.01 | 0.01 | 0.02 | 0.01 | 0 | 0.02 |
| 26 | Southern Great Escarpment | Present | 0.01 | 0 | 0.02 | 0 | -0.01 | 0 | -0.01 | -0.02 | 0.01 |
| 27 | Madagascar* | Absent | 0 | 0 | 0 | 0 | 0 | 0 | -0.03 | -0.03 | -0.03 |
| 28 | Caucasus | Absent | 0 | 0 | 0.01 | 0 | -0.01 | 0 | -0.02 | -0.03 | -0.02 |
| 29 | Zagros Mountains | Absent | 0 | -0.01 | 0 | -0.01 | -0.01 | 0 | -0.03 | -0.03 | -0.03 |
| 30 | Urals | Absent | -0.02 | -0.02 | -0.02 | -0.01 | -0.01 | -0.01 | -0.05 | -0.05 | -0.04 |
| 31 | Russia | Absent | 0 | -0.01 | 0.01 | -0.01 | -0.02 | -0.01 | -0.07 | -0.07 | -0.07 |
| 32 | Hindukush-Himalaya* | Present | 0.07 | 0.06 | 0.08 | 0.02 | 0.01 | 0.03 | -0.07 | -0.07 | -0.06 |
| 33 | India* | Present | 0.01 | 0 | 0.01 | -0.03 | -0.03 | -0.03 | -0.02 | -0.03 | 0 |
| 34 | Japan | Absent | 0 | 0 | 0.01 | -0.01 | -0.01 | -0.01 | -0.02 | -0.02 | -0.02 |
| 35 | Korea | Absent | 0 | -0.01 | 0.01 | 0 | 0 | 0 | -0.03 | -0.03 | -0.03 |
| 36 | Taiwan* | Absent | 0.01 | 0.01 | 0.01 | 0 | 0 | 0.01 | -0.01 | -0.01 | 0 |
| 37 | China | Present | 0.06 | 0.05 | 0.07 | 0.01 | 0.01 | 0.02 | -0.03 | -0.03 | -0.03 |
| 38 | Philippines* | Present | 0.02 | 0.01 | 0.03 | 0.01 | 0 | 0.01 | -0.03 | -0.04 | -0.03 |
| 39 | Southeast Asia* | Present | 0.04 | 0.02 | 0.06 | 0 | -0.01 | 0.01 | -0.07 | -0.08 | -0.07 |
| 40 | Sumatran Islands* | Present | 0.02 | 0.01 | 0.03 | -0.02 | -0.02 | -0.02 | -0.01 | -0.02 | 0.01 |
| 41 | Kapuas Mountains* | Present | 0.01 | -0.01 | 0.02 | 0.01 | 0 | 0.01 | -0.01 | -0.02 | 0 |
| 42 | Sulawesi* | Present | 0.01 | 0.01 | 0.01 | 0 | 0 | 0 | 0 | -0.01 | 0.01 |
| 43 | Papua New Guinea* | Present | 0.01 | 0 | 0.03 | 0 | -0.01 | 0.01 | -0.05 | -0.06 | -0.05 |
| 44 | Central Australia | Absent | -0.06 | -0.06 | -0.05 | -0.03 | -0.03 | -0.03 | -0.05 | -0.05 | -0.04 |
| 45 | Great Dividing Range* | Absent | -0.08 | -0.08 | -0.07 | -0.06 | -0.07 | -0.06 | -0.09 | -0.09 | -0.08 |

| | | | | | | | | | | | |
|---|---|---|---|---|---|---|---|---|---|---|---|
| 46 | Southern Alps New Zealand | Absent | 0 | -0.01 | 0 | -0.01 | -0.01 | -0.01 | -0.03 | -0.04 | -0.03 |


1. B. G. Freeman, M. Strimas-Mackey, E. T. Miller, Interspecific competition limits bird species' ranges in tropical mountains. *Science* **377**, 416–420 (2022).

2. I. Quintero, W. Jetz, Global elevational diversity and diversification of birds. *Nature* **555**, 246–250 (2018).

3. R Core Team, *R: A Language and Environment for Statistical Computing* (R Foundation for Statistical Computing, 2022).

4. J. A. Tobias, *et al.*, AVONET: morphological, ecological and geographical data for all birds. *Ecology Letters* **25**, 581–597 (2022).


**Data sources:**

Hyperlinks to these sources are below:

1. I. Quintero, W. Jetz, Global elevational diversity and diversification of birds. *Nature* **555**, 246–250 (2018): Supplementary Table 1 at: https://www.nature.com/articles/nature25794#Sec20

2. J. A. Tobias, *et al.*, AVONET: morphological, ecological and geographical data for all birds. *Ecology Letters* **25**, 581–597 (2022): AVONET Supplementary dataset 1.xlsx at: https://figshare.com/s/b990722d72a26b5bfead?file=34480856

3. B. G. Freeman, M. Strimas-Mackey, E. T. Miller, Interspecific competition limits bird species' ranges in tropical mountains. *Science* **377**, 416–420 (2022): https://zenodo.org/record/6450245#.Y_DIAxxByV4
